# Single-cell immune repertoire and transcriptome sequencing reveals that clonally expanded and transcriptionally distinct lymphocytes populate the aged central nervous system in mice

**DOI:** 10.1101/2020.05.04.077081

**Authors:** Alexander Yermanos, Daniel Neumeier, Ioana Sandu, Mariana Borsa, Ann Cathrin Waindok, Doron Merkler, Annette Oxenius, Sai T. Reddy

## Abstract

Neuroinflammation plays a crucial role during ageing and various neurological conditions, including Alzheimer’s disease, multiple sclerosis and infection. Technical limitations, however, have prevented an integrative analysis of how lymphocyte immune receptor repertoires and their accompanying transcriptional states change with age in the central nervous system (CNS). Here, we leveraged single-cell sequencing to simultaneously profile B cell receptor (BCR) and T cell receptor (TCR) repertoires and accompanying gene expression profiles in young and old mouse brains. We observed the presence of clonally expanded B and T cells in the central nervous system (CNS) of aged mice. Furthermore, many of these B cells were of the IgM and IgD isotype and had low levels of somatic hypermutation. Integrating gene expression information additionally revealed distinct transcriptional profiles of these clonally expanded lymphocytes. Our findings implicate that clonally related T and B cells in the CNS of elderly mice may contribute to neuroinflammation accompanying homeostatic ageing.

## Introduction

Ageing of the immune system, commonly referred to as immune senescence, has been shown to hamper adaptive immune responses in the context of vaccination, viral infection and neurological conditions ^1–4^. Host-mediated immunopathology is sometimes increased in the elderly during immune challenges ^5^, further highlighting the importance of studying adaptive immunity in the context of ageing. A number of phenotypic and functional alterations have been linked to dysfunctional adaptive immunity, including the diminished production of new naïve B and T cells ^6–8^, increased host-immunopathology during viral infections and disease ^5,9^, increased exhaustion and elevated expression levels of dysfunction markers ^10,11^. A prominent example revealed that CD8 T cells up-regulated classical exhaustion markers (PD-1, LAG3, 2B4, CD160) when comparing 4-month-old to 12-month-old mice. Strikingly, the difference between 12-month-old and 18-month-old mice demonstrated an even more dramatic increase in the aforementioned exhaustion markers, exemplified by an even larger increase in PD-1^+^ naïve T cells in 18-month-old mice. A comparable population of age-associated B cells has been recently described and characterized by absence or presence of specific surface markers: CD21^−^, CD23^−^, Tbet^+^, and CD11c^+ 12–14^. This population increases with age and has been associated with the production of autoantibodies and pro-inflammatory cytokines, despite the antigen-specificity of these B cells remaining unknown.

A recent study using high-dimensional mass-cytometry reported an increased percentage of B and T cells located in the elderly murine CNS ^15^. However, the phenotypic analysis of this study was primarily focused on myeloid cells and microglia, thereby leaving the phenotype of adaptive immune cells uncharacterized. The importance of adaptive immune cells in the context of neurological disorders has been further highlighted by the success of monoclonal antibody therapies targeting cells of the lymphocyte branch ^16,17^. In one instance, a B cell-depleting anti-CD20 monoclonal antibody was used for the treatment of multiple sclerosis ^18–20^. These findings have implicated an antibody-independent role of adaptive immune cells in CNS pathology, as the primary antibody producing cells (plasma cells) do not express CD20 ^21–23^. Furthermore, the aged CNS is populated with adaptive immune cells in the context of various diseases ^24,25^, thereby warranting further investigation into CNS-resident lymphocytes. In particular, as the prevalence of neurological diseases tends to increase with age ^26–28^, elucidating key features of adaptive immunity (i.e., lymphocyte clonal diversity and clonal expansion along with defined transcriptional states) in aged brain tissue will be of utmost importance. Due to technological limitations however, previous studies were unable to provide such information, thereby ignoring crucial features of adaptive immunity in the CNS of aged individuals.

Advances in deep sequencing and microfluidic-based technologies have revolutionized the resolution with which we are now able to extract biologically relevant information from adaptive immune receptor repertoires (immune repertoires) ^29–31^. Immune repertoires represent a vast and diverse collection of B and T cell receptors (BCR and TCR, respectively) and can be used to quantify both past and ongoing immune responses of an individual. The majority of studies in the past have focused primarily on delineating repertoire metrics based on a single variable region: the variable heavy (V_H_) chain in case of BCRs ^32,33^ or the variable beta (Vβ) chain in case of TCRs ^33–35^. However, new single-cell sequencing technologies are now enabling a more comprehensive characterization of immune repertoires by capturing both BCR variable light (V_L_) and V_H_ regions and TCR variable alpha (Vα) and Vα chains ^36,37^. For example recent studies employing single-cell TCR sequencing have discovered convergent sequence motifs correlating to MHC-peptide antigens from bacterial or viral pathogens ^38–40^. Single-cell sequencing can also recover transcriptional profiles, which can be linked to immune repertoire clonal identifiers to thus obtain a comprehensive phenotypic characterization of lymphocytes ^36^. For example, single-cell profiling of BCR repertoires and transcriptomes has recently revealed a high degree of bystander activation during influenza vaccination ^36^. This combined immune repertoire and transcriptome profiling has also been employed to obtain unprecedented resolution of lymphocyte dynamics in the context of cancer ^41,42^.

Here, we performed single-cell immune repertoire and transcriptome sequencing of B and T cells from 3-month-old, 12-month-old, and 18-month-old mice in order to phenotypically characterize and determine the extent of clonal expansion of adaptive immune cells in the aged CNS. Our strategy to profile B and T lymphocytes allowed us to relate repertoire parameters such as clonal expansion, germline gene usage and isotype (in case of B cells) to gene expression profiles at the single-cell resolution. Our findings demonstrated the presence of highly expanded B and T cell clonal lineages in the aged CNS. We could additionally demonstrate that expanded B cells in the aged CNS were predominantly of the IgM isotype and exhibit low levels of somatic hypermutation. While we observed that clonally expanded CNS lymphocytes had distinct transcriptional profiles compared to unexpanded clones, there were few age-associated changes in gene expression. Taken together, our findings demonstrate the accumulation of clonally expanded B and T lymphocytes in the CNS upon ageing that may contribute to age-associated neuroinflammation.

## Results

### Single-cell sequencing of immune repertoires and transcriptomes of lymphocytes from murine CNS

To investigate the lymphocytes in the aged CNS, we performed single-cell sequencing on B and T cells isolated from entire brains of pooled 3-month-old (n=4), 12-month-old (n=4) and 18-month-old C57/BL6j male mice (n=4). We additionally processed and sequenced the brain of an individual 18-month-old mouse to ensure both reproducibility and to allow the detection of common immune cell clones shared between mice. B and T lymphocytes were isolated by performing fluorescence-activated cell sorting (FACS) of cell suspensions derived from whole brains of each cohort. B cells were isolated based on CD19^+^ B220^+^ surface expression, whereas T cells were sorted based on surface expression of CD3^+^ and CD4^+^ or CD8^+^ (Figure 1A). Following sorting, the entirety of the three isolated immune cell populations were pooled and supplied as input for single-cell sequencing using the 10X Genomics Chromium pipeline. Following cell lysis, reverse transcription, and introduction of cell-specific nucleotide barcodes, we split each barcoded sample into three to prepare gene expression libraries (GEX), T cell V(D)J libraries, or B cell V(D)J libraries separately.

**Figure 1.**
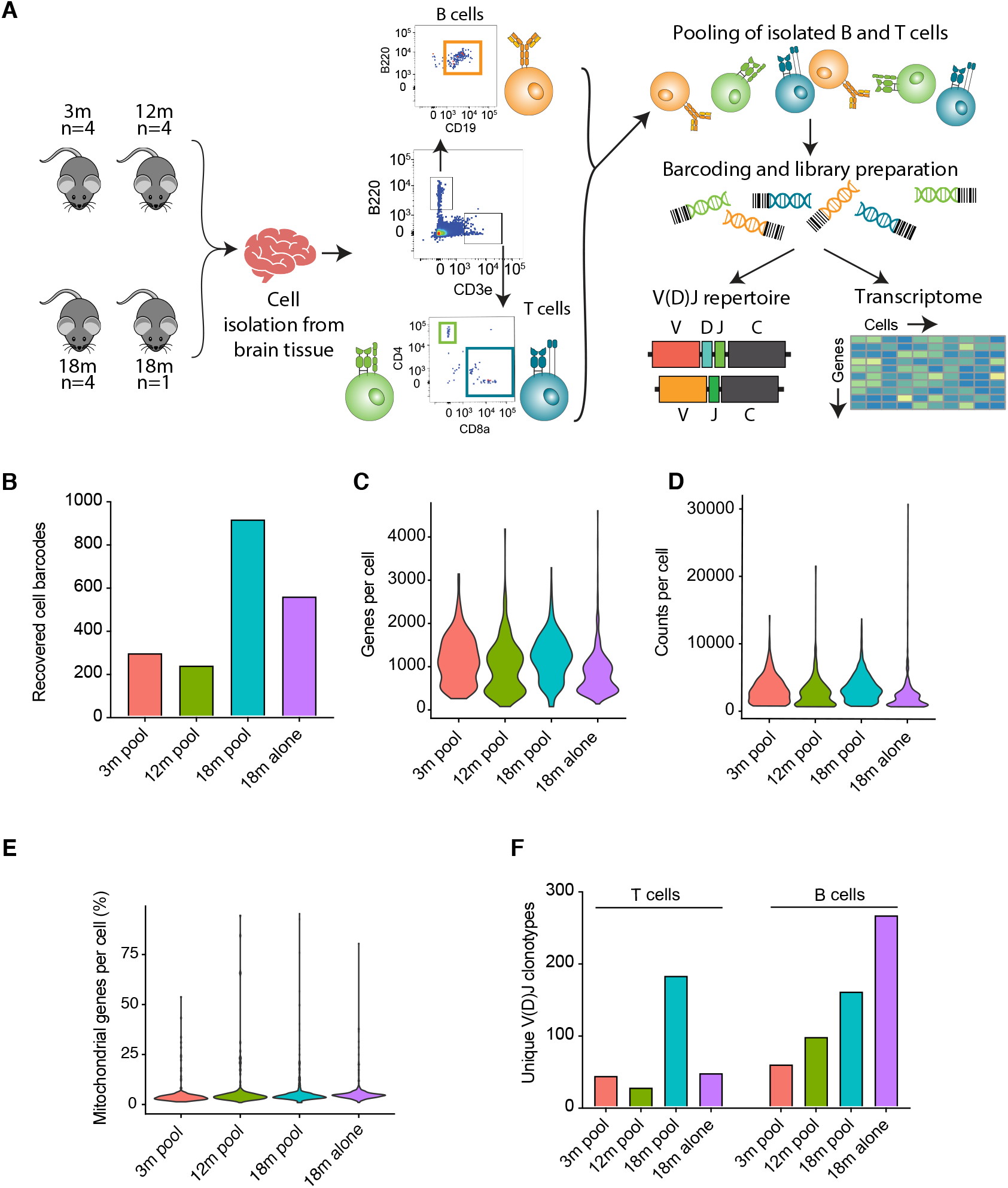
Single-cell sequencing recovers repertoire and gene expression information of B and T cells from the aged murine central nervous system. (A) Experimental overview of isolation of B and T cells from entire murine brains. (B) Number of recovered cell barcodes following Single-cell sequencing of gene expression libraries for each of the experimental cohorts. (C) Unique genes detect per cell for each of the experimental cohorts. (D) Total number of RNA molecules detected within each cell. (E) Percentage of mitochondrial genes for each cell. (F) Number of unique clones from V(D)J sequencing library for each of the experimental cohorts. Clone is defined as identical, paired amino acid CDR3 sequences (CDRβ3+CDRα3 for T cells, CDRH3+CDRL3 for B cells) with exactly one heavy chain and one light chain.

Following deep-sequencing of the three libraries per cohort, we were able to recover transcriptional information for approximately two thousand distinct cells (Figure 1B). In accordance with previous reports of increased B and T cells in the aged mouse brain ^15^, we observed a higher number of cells from both cohorts of 18-month-old animals (Figure 1B). We did not observe an increased number of recovered cells in the pooled 18-month-old relative to the individually processed 18-month-old brain, suggesting potential cell loss during pooling and sequencing library preparation. Nevertheless, we detected on average ~1000 genes per cell across all cohorts, with an average of approximately 2700 counts per cell (Figures 1C, 1D). We additionally observed that, on average, fewer than 5% of counts per cell mapped to mitochondrial genes, implying adequate sequencing quality (Figure 1E).

We next analyzed full-length paired variable region clones (V_H_:V_L_ for B cells and Vβ:Vα for T cells); a clone was defined based on the amino acid sequence of complementary determining region 3 (CDR3) of both variable regions (i.e. CDRH3+CDRL3 for BCR, CDRβ3+CDRα3 for TCR). We observed increased clonal diversity for both B and T cells in the aged cohorts (Figure 1F), as well as for the BCR repertoire from the individual 18-month-old mouse compared to the pooled 18-month-old cohort (Figure 1F), potentially suggesting mouse-heterogeneity with regards to clonal diversity or that technical biases arose during sample pooling.

### Clonally expanded B and T cells are located in the murine CNS of elderly mice

After witnessing the clonal diversity in the immune repertoires of aged mice, we next questioned whether this was due to random infiltration of unrelated, circulating lymphocytes or due to infiltration of clonally related B and T cells. V(D)J recombination generates a diverse panel of BCRs and TCRs, such that they are essentially unique to an individual recombination event ^43^. Thus, finding two or more lymphocytes with identical clonal sequences (CDRH3+CDRL3 or CDRβ3+CDRα3) strongly supports that clonal expansion occurred. We therefore quantified the number of cell barcodes for the top 40 most expanded B and T cells for the four cohorts (Figures 2A, 2B). In the 3-month-old pooled brain, we observed only 7 BCR clones corresponding to two distinct B cells (Figure 2A). Comparing this to the other two cohorts, revealed an increase in B cell clonal expansion for both 12-month-old and 18-month-old mice (Figure 2A). Due to our experimental setting, it is not possible to resolve whether this clonal expansion is due to BCR clones recovered in multiple mice (public clones) or clonal expansion within a single mouse. To further investigate this, we performed a similar quantification for the individually processed 18-month-old mouse, which revealed immense clonal expansion relative to the low number of recovered cells (Figures 2A, 1B). For example, 33 cells were present in the most expanded clonal family, and additionally there were still 3 cells corresponding to the 40th most expanded clone (Figure 2A). The definition of a clonal family here is based on exact CDRH3+CDRL3 amino acid sequences, which thus excludes BCR sequences that have somatic hypermutation in the CDR3 regions. Thus, we performed an additional clonotyping analysis based on identical V_H_ and V_L_ germline genes and identical CDRH3 and CDRL3 amino acid sequence lengths, which resulted in only a difference for three clones (Figure S1), thereby confirming that our initial clonal family definition sufficiently described clonal expansion.

**Figure 2.**
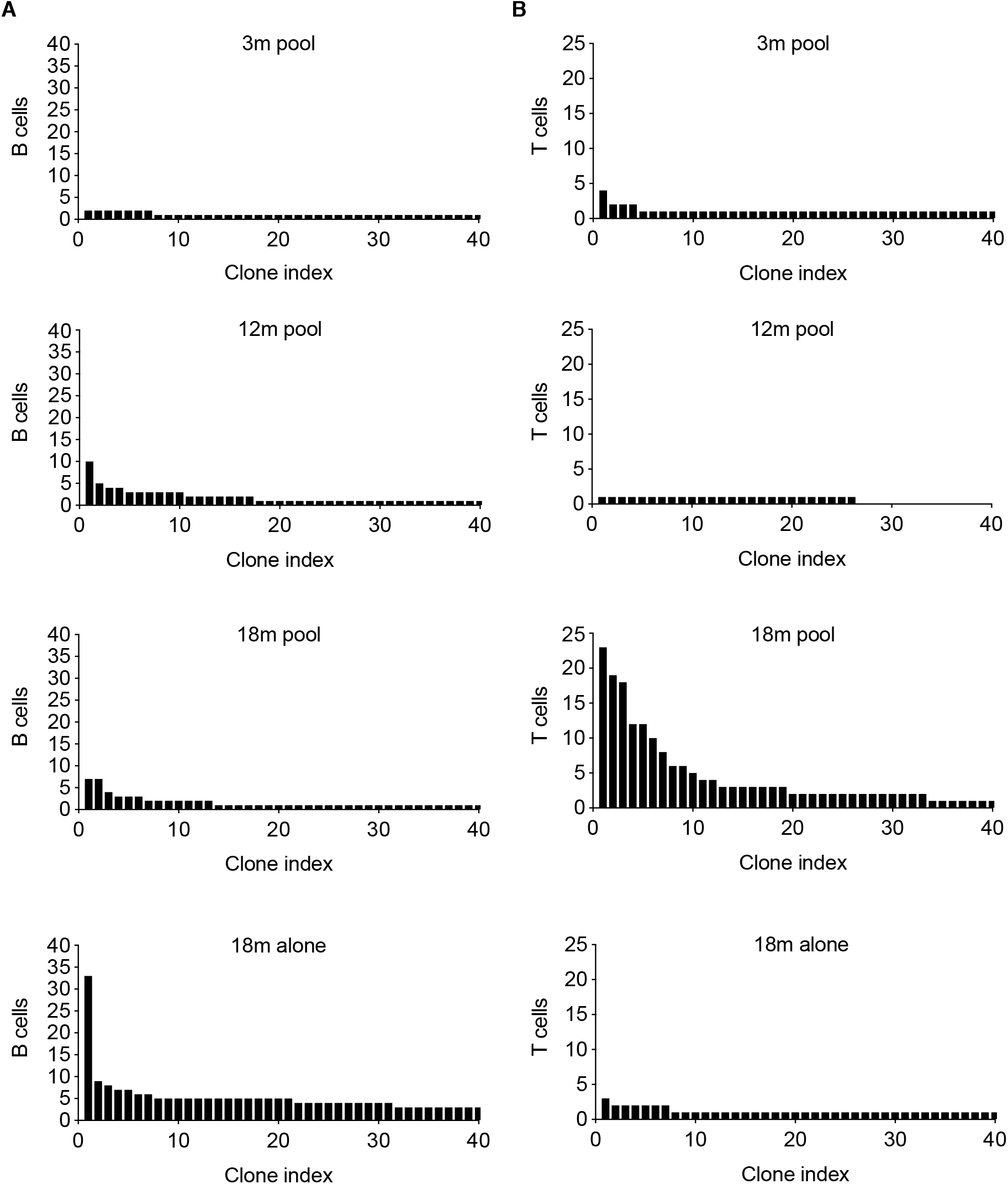
Clonally expanded B and T cells in the aged CNS. (A-B) The number of recovered cells corresponding to the top 40 most expanded clones for B and T cell repertoires, respectively. Clone is defined as identical, paired amino acid CDR3 sequences (CDRβ3+CDRα3 for T cells, CDRH3+CDRL3 for B cells) with exactly one heavy chain and one light chain. Clone index corresponds to the clonal rank determined by frequency.

The presence of clonally expanded B cells in the aged CNS prompted us to question whether similar expansion profiles would be observed for T cells. We therefore performed the identical analysis for the TCR repertoires for each age cohort. Measuring the clonal expansion for the top 40 T cell clones demonstrated that we could again recover multiple cells containing the same clonal sequence (Figure 2B). In the 3-month-old and 12-month-old pooled brain TCR repertoires, there were very few clonal sequences corresponding to more than 2 cells (Figure 2B). However, in the pooled 18-month-old cohort, clonal expansion was nevertheless detected, with the most expanded clone corresponding to 23 cells. While it is again not possible to entirely delineate whether these are arising from a single mouse or public clones, combining these results with the individually sequenced 18-month-old mouse nevertheless supports that clonally expanded lymphocytes are present in the aged CNS (Figure 2B).

### BCR repertoires in aged CNS have characteristics of naïve B cells

After observing clonal expansion in the brains of aged mice, we next investigated whether these B cells had undergone class-switch recombination, which would be a correlate of a previous interaction with cognate antigen. We therefore extracted the heavy chain isotype for each B cell clonal lineage across all four age cohorts. For clonal families containing more than one distinct cell, we used the isotype corresponding to the majority of cells within that family. The majority of the expanded clones were of the IgM isotype for all age cohorts, although surprisingly there were many B cell clones corresponding to the IgD isotype for all four cohorts (Figure 3A). We next determined the isotype distribution of clonally expanded B cells (Figure 2A), which revealed that the majority of clonally expanded families contained cells of either the IgM, IgD or a combination of the two (Figure 3B). The unexpected occurrence of exclusive IgD clones was also observed as trend in many B cells of aged mice (Figure 3C). We further assessed whether these BCR sequences were supported by multiple unique molecular identifiers (UMIs), because these isotype results could have been a result of not sampling enough mRNA molecules for each cell, thereby not detecting co-expressed IgM. However, we observed that almost all (>97%) of these IgD clones were inferred using multiple UMIs (Figure 3D), suggesting sufficient sequencing and sampling depth and thereby excluding to have missed co-expression of IgM.

**Figure 3.**
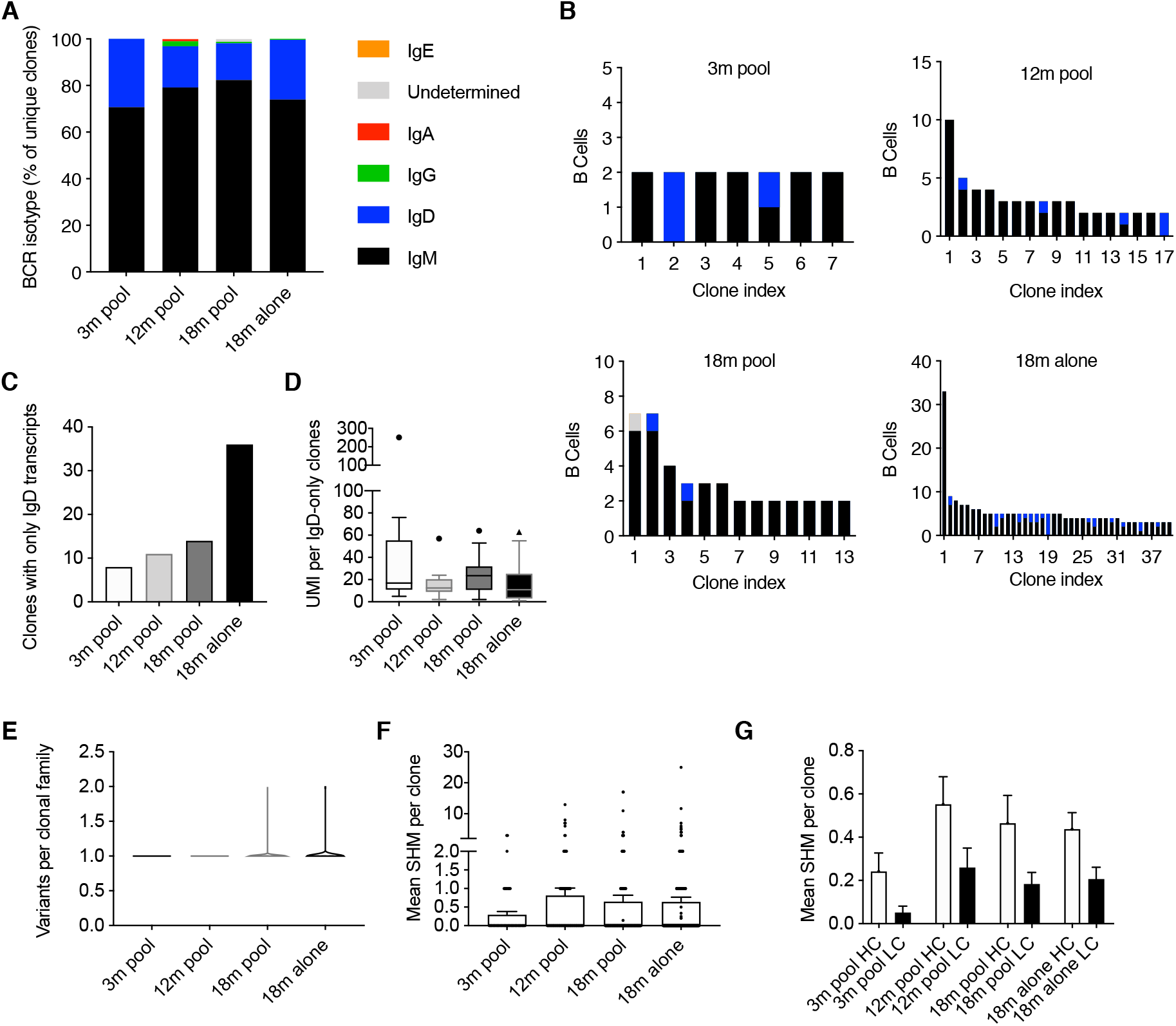
B cells in the CNS are primarily of the IgM isotype with minor somatic hypermutation. (A) Percentage of unique clones with a given BCR. For those clones with multiple cells, the majority isotype was selected. (B) Isotype distribution for those expanded (>1 cell barcode) B cell clones. (C) Number of clones with exclusively IgD BCR transcripts. (D) Tukey box-plot displaying the number of unique molecular identifiers (UMI) per those clones with reads mapping exclusively to the IgD isotype. Horizontal line within the box indicates the median and dot above. (E) The number of somatic variants per expanded B cell clone. Variant is defined as unique, combined V_H_ + V_L_ amino acid sequence. (F) The mean number of nucleotide somatic hypermutations (SHMs) per B cell clone in the full-length V and J regions across both heavy (HC) and light chain (LC). (G) The mean number of nucleotide substitutions per B cell clone detected on either the HC or light chain LC. Both HC and LC sequences correspond to full-length V and J regions.

Our previous analyses demonstrating that relaxing clonotyping stringency did not alter clonality in BCR repertoires, suggested that there were not substantial amino acid mutations in the CDR3 regions of B cells in the CNS. This was further implicated by the lack of class-switched B cells. We therefore next investigated whether a similar lack of somatic hypermutation was observed in the remaining regions of the BCRs. To accomplish this, we first quantified the number of somatic variants (defined by unique, full-length, paired V_H_-V_L_ nucleotide sequences) within each clonal family. This demonstrated that the majority of these clonal families containing more than one cell had only one unique antibody sequence (i.e. all cells in a clonal family produced the same antibody) for all age cohorts (Figure 3E). We next determined divergence from germline for each clonal family which showed that despite low mutation levels for all BCR repertoires, there was a trend of increased somatic hypermutation in the aged cohort, with certain clones diverging from the reference germline by more than 10 nucleotide mutations (Figure 3F). Single-cell sequencing allows us to separately analyze V_H_ and V_L_ regions, thus we additionally determined whether somatic hypermutation was distributed evenly across the two regions. In line with previous findings, the V_H_ region was the dominant target for somatic hypermutation across all cohorts (Figure 3G). Taken in concert, these results suggest that the majority of B cells express IgM or IgD BCRs and that there is a slight increase in somatic hypermutation in BCR repertoires of aged mice.

### Assessing sequence similarity of immune repertoires

After observing comparable isotype distributions in the BCR repertoires of young and old mice, we next questioned whether we could detect hints of clonal convergence using other repertoire metrics. While multiple bioinformatics analyses exist to quantify the similarity between repertoires, we assessed whether there were identical BCR or TCR clones (defined as identical amino acid sequences of CDRH3+CDRL3 for BCR and CDRβ3+CDRα3 for TCR) shared between the different age groups. This analysis revealed that for both BCR and TCR there were no identical clones found between the different age groups (Figure 4A). However, when restricting our analysis to a single variable chain, (clone defined as identical amino acid sequence of CDRH3, CDRβ3, CDRL3 or CDRα3), we observed substantial clonal overlap in variable light chains of 18-month-old cohorts (Figure 4A). We next determined if relaxing the definition of clonal overlap would alter metrics of repertoire similarity between the aged cohorts. To address this, we employed metrics from graph theory to quantify sequence similarity between clones both within and across repertoires ^29^. In sequence similarity networks, clones are represented as nodes and are connected via edges to other clones with high sequence similarity (four or less amino acid mutations across the paired CDR3s, Figure 4B, 4C). Information corresponding to age and clonal frequency is demonstrated by color and size of nodes, respectively. We first illustrated the overall connectivity between all repertoires by including each unique clone in our similarity network. We additionally created similarity networks to demonstrate the relationship between the connected nodes, which clearly demonstrated a higher degree of sequence similarity between BCRs relative to TCRs across cohorts (Figure 4B, 4C). There were several clusters in the BCR sequence similarity network in which only clones from the CNS repertoires of aged mice were connected (Figure 4C), potentially suggesting these clones may have a similar antigen-specificity and could accumulate with age. Finally, we quantified the percentage of unique clones using each V germline gene for the BCR and TCR variable chains for all cohorts and subsequently performed hierarchical clustering. This revealed that the 18-month-old cohorts clustered together for BCR V_H_ and V_L_ and TCR Vα chains, whereas the pooled repertoires for 12-month-old and 18-month-old mice clustered together for the TCR Vα chain (Figures 4D). Together, these findings suggest that V-gene usage similarity of immune repertoires increases in the elderly CNS.

**Figure 4.**
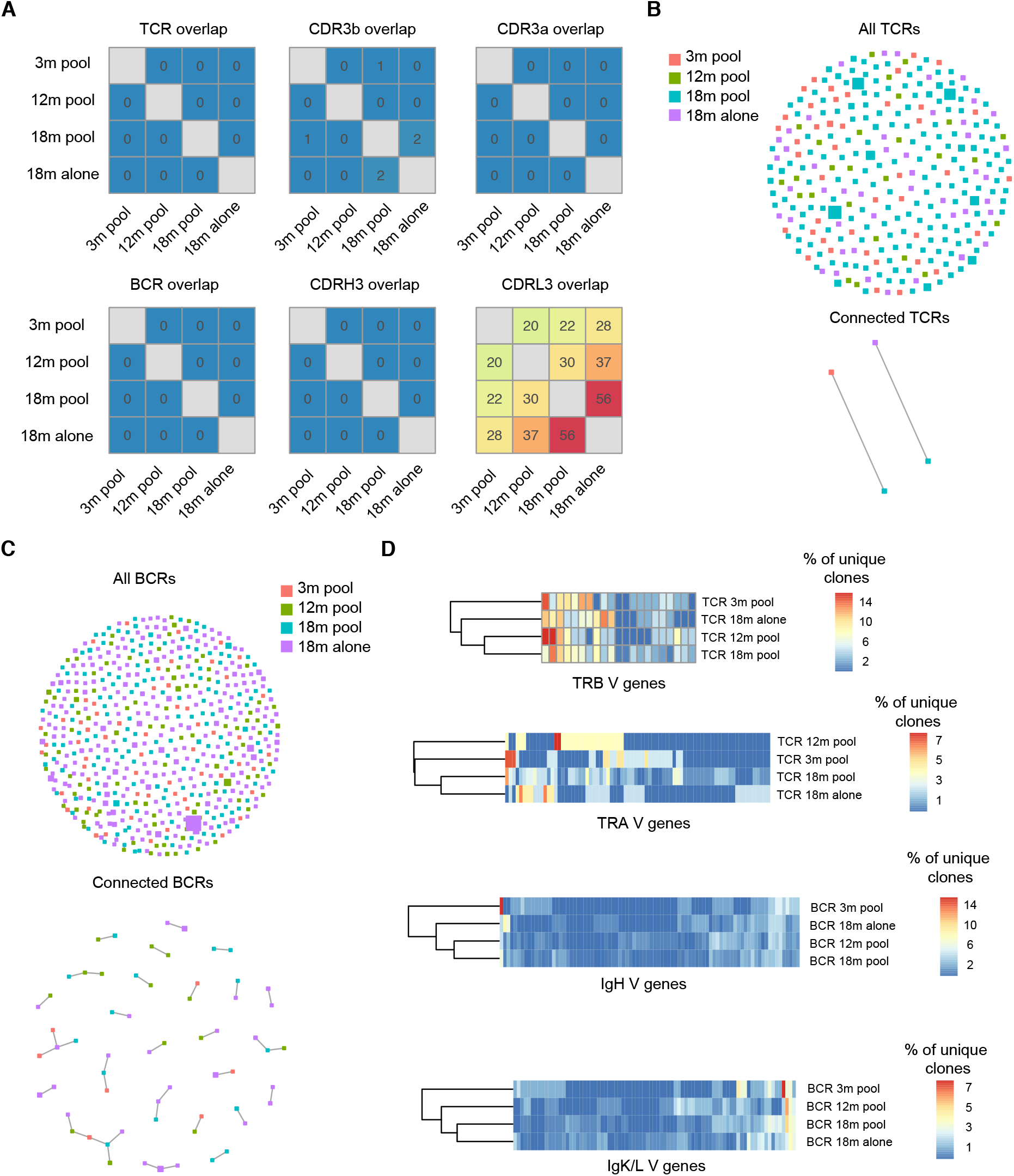
Minor clonal convergence between CNS immune repertoires. (A) Heatmap depicting clonal overlap between repertoires. Clonal overlap is defined as the number of identical amino acid CDR3 sequences shared between repertoires. BCR and TCR overlap corresponds to the paired CDRH3+CDRL3 / CDRβ3+CDRα3 amino acid sequence. (B-C) Similarity networks depicting clonal relatedness and expansion between T cell and B cell repertoires, respectively. Size of nodes corresponds to relative clonal expansion. Color of node corresponds to experimental cohort. Edges between nodes indicate two clones (paired CDRH3+CDRL3 / CDRβ3+CDRα3 amino acid sequence) that are separated by four or less amino acid mutations. Lower panel highlights those clones containing edges, whereas upper panel depicts all clones. (D) Percent of unique clones using each V gene for each experimental cohort.

### Detection of transcriptionally distinct B and T cell clusters

Since single-cell sequencing workflows also enable parallel interrogation of transcriptomes, we are able to assess the gene expression profiles of these of B and T cells in the CNS of the various aged cohorts. We used transcriptome sequencing data to perform unsupervised clustering and uniform manifold approximation projection (UMAP) to group cells with similar gene expression profiles. This resulted in 11 distinct clusters which were populated by cells from all cohorts (Figures 5A, 5B). We next determined whether single-cell transcriptome data was sensitive to B and T cell clusters, as determined by cell-surface markers used during the FACS isolation. Overlaying the expression levels of CD19, CD3e, CD8a and CD4 demonstrated that while we did observe populations corresponding to these cell types, there were also clusters detected that did not express these markers (Figure 5C). For example, microglial genes (Trem119 and OLFML3) were detected in cluster 4, indicating possible impurities arising from FACS isolation (Figure 5D). Quantifying the number and percentage of cells found in each cluster indicated that cells from all age cohorts populated the clusters containing B cell genes (1,3,5,7,8 based on CD79, CD19, CD20) and T cell genes (clusters 0,2,9 based on CD3e, CD8a, CD4) (Figure 5D). Taken in concert, these analyses revealed comparable transcriptional profiles across all age cohorts and the need to focus our transcriptional analysis on those specific B and T cell lymphocytes detected in our V(D)J sequencing library.

**Figure 5.**
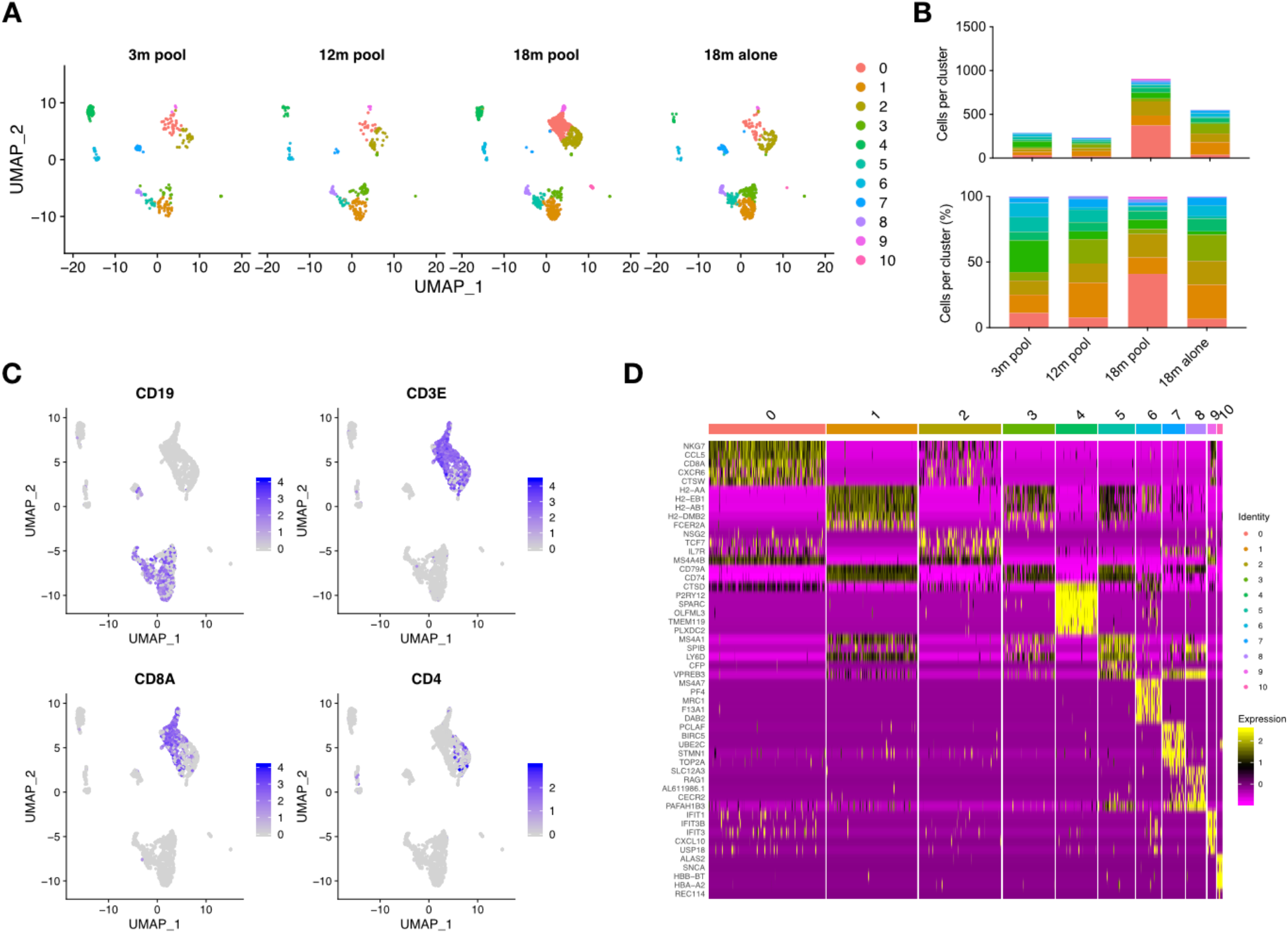
Gene expression libraries reveal transcriptional similarities between young and old CNS. (A) Unsupervised clustering and uniform manifold approximation projection (UMAP) reveals distinct transcriptional clusters in the CNS. Each dot represents a cell recovered in the gene expression library. Colors correspond to transcriptional cluster membership. (B) The number and percent of cells found within each cluster for each of the experimental cohorts. (C) Expression for CD19, CD3e, CD8a, and CD4 for all cells recovered in the gene expression libraries. Intensity corresponds to normalized expression. Each dot represents a cell. (D) Heatmap displaying clustering-defining genes for the 11 clusters. Cells were randomly sampled from within each cluster proportional to cluster’s total number of cells. Intensity corresponds to normalized gene expression.

### Profiling transcriptional landscape in aged CNS lymphocytes

After observing clusters corresponding to cells other than lymphocytes, we next determined whether we could restrict our transcriptome analysis by only including those barcodes detected in the V(D)J libraries. Since the 10X Genomics single-cell sequencing workflow consists of labeling each cell with unique barcodes before separating cDNA as input into immune repertoire or gene expression libraries, it was possible to selectively investigate cells for which both repertoire and transcriptome data was available (n=1008 cells). Highlighting cells for which full-length variable region repertoire information was available demonstrated that our previously hypothesized clusters (Figure 5) were indeed populated with B and T cells (Figure S2A). Consistent with our previous results (Figure 5A), the B and T cells from all age cohorts colocalized based on gene expression information, further highlighting similarities between the aged cohorts when using an unbiased clustering approach (Figures S2B, S2C). Next we investigated if separating B and T cells would reveal any age-associated transcriptional changes between cohorts. By restricting phenotypic analysis to cells for which we had complete BCR or TCR sequence information (paired CDR3s, full-length V(D)J sequence) and performing unbiased clustering and UMAP resulted in 5 and 4 transcriptional clusters for the T and B cells, respectively (Figures 6A). We again observed comparable frequencies of cluster membership across all age cohorts (Figures 6B), further suggesting that lymphocytes in the aged CNS maintain comparable transcriptional profiles to those in young mice.

**Figure 6.**
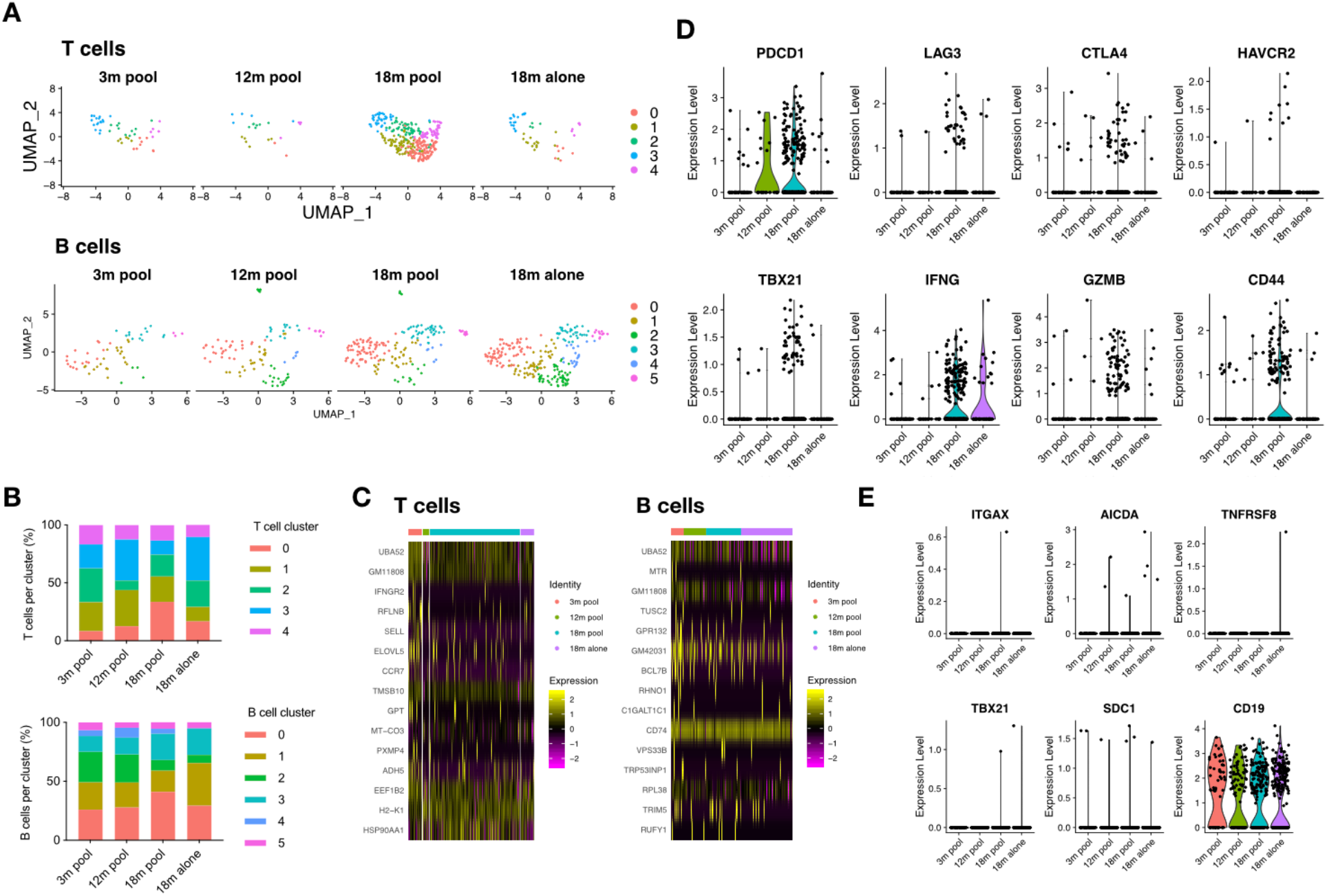
Transcriptional signatures of clonally expanded lymphocytes. (A) Unsupervised clustering and uniform manifold approximation projection (UMAP) was performed separately on either B or T cells that were also detected in V(D)J libraries. Each dot represents a single cell. Color corresponds to transcriptional cluster specific to T or B cell analysis. (B) The percent of T (top) or B (bottom) cells found within each cluster for each of the experimental cohorts. Clusters were determined using either only T cells or only B cells found in V(D)J libraries. (C) Heatmap displaying clustering-defining genes for T and B cell clusters. Clusters were calculated separately for B cells and T cells. Cells were randomly sampled from within each cluster proportional to cluster’s total number of cells. Intensity corresponds to normalized gene expression. (D) Normalized gene expression of for T cells for selected genes. (E) Normalized gene expression of for B cells for selected genes.

We next determined if any age-associated differences in the transcriptional profiles of lymphocytes was observed when cluster membership was not taken into account. To this end we computed the differential expression between the cohorts and found minor differences (Figure 6C). However, we observed a trend of upregulation of markers associated with T cell function or exhaustion such as PD-1, Lag3, CTLA4, Tim3 (HAVCR2) in aged but not young mice (Figure 6D). Other genes relevant to T cell function, such as expression of granzymes and interferon-gamma followed a similar trend of upregulation in the 18-month-old animals, albeit with a low number of cells (Figure 6D). With the exception of CD19, characteristic B cell markers, including age-associated genes (CD11c and TBX21) and differentiation genes (AID, CD138, CD30) showed no major differences across cohorts (Figure 6E).

### Clonally expanded lymphocytes have distinct transcriptional profiles

Since we could observe clonal expansion in immune repertoires aged CNS, we investigated the transcriptional landscape of these cells. Highlighting expanded clones (clonal families corresponding to at least two distinct cell barcodes) suggested transcriptional-dependent clustering for both B and T cells (Figures 7A, 7B). While expanded T cells were distributed across all clusters (excluding cluster 3), we observed that the majority were CD8 T cells (Figures 7C), clarifying our previous clonal expansion findings (Figure 2B). In order to detect transcriptional signatures, we explicitly quantified differential gene expression between clonally expanded lineages and those containing only a single cell. This comparison between expanded to unexpanded cells resulted in 26 and 57 genes differentially regulated (p.adj < 0.01) for T and B cells, respectively (Figures 7D, 7E). Strictly taking the top genes based on *p*-values demonstrated that expanded T cells expressed higher levels of NKG7, CCL5, CXCR6, CCL4, Granzyme K, and ID2 and had downregulated markers associated with a central memory phenotype, such as CD62L, CCR7, LEF1, and IL7R (Figure 7D). The phenotypic profile of expanded B cells was less clear, although some relevant genes such as CCR7 and ones involving proliferation (e.g. BTG1) were detected as downregulated in expanded cells (Figures 7E). Quantifying gene expression for a panel of relevant genes demonstrated a trend that clonally expanded T cells expressed more PD-1 (PDCD1), interferon gamma (IFNG), and granzyme B (GZMB) (Figure S3A), suggesting recent or persistent activation. Performing a similar analysis using a panel of age- and activation-associated B cell markers resulted in minor differences in gene expression between expanded and unexpanded B cells, although this analysis may have been limited by low count numbers for the selected genes (Figure S3B). Extending our analysis to quantify differences between the expanded cells of aged versus young CNS lymphocytes did not reveal major changes in transcription, again consistent with our previous findings (Figures S4A, S4B, S4C). Taken together, these results demonstrate distinct transcriptional phenotypes of clonally expanded lymphocytes in the CNS that are robust to age-associated changes following homeostatic ageing.

**Figure 7.**
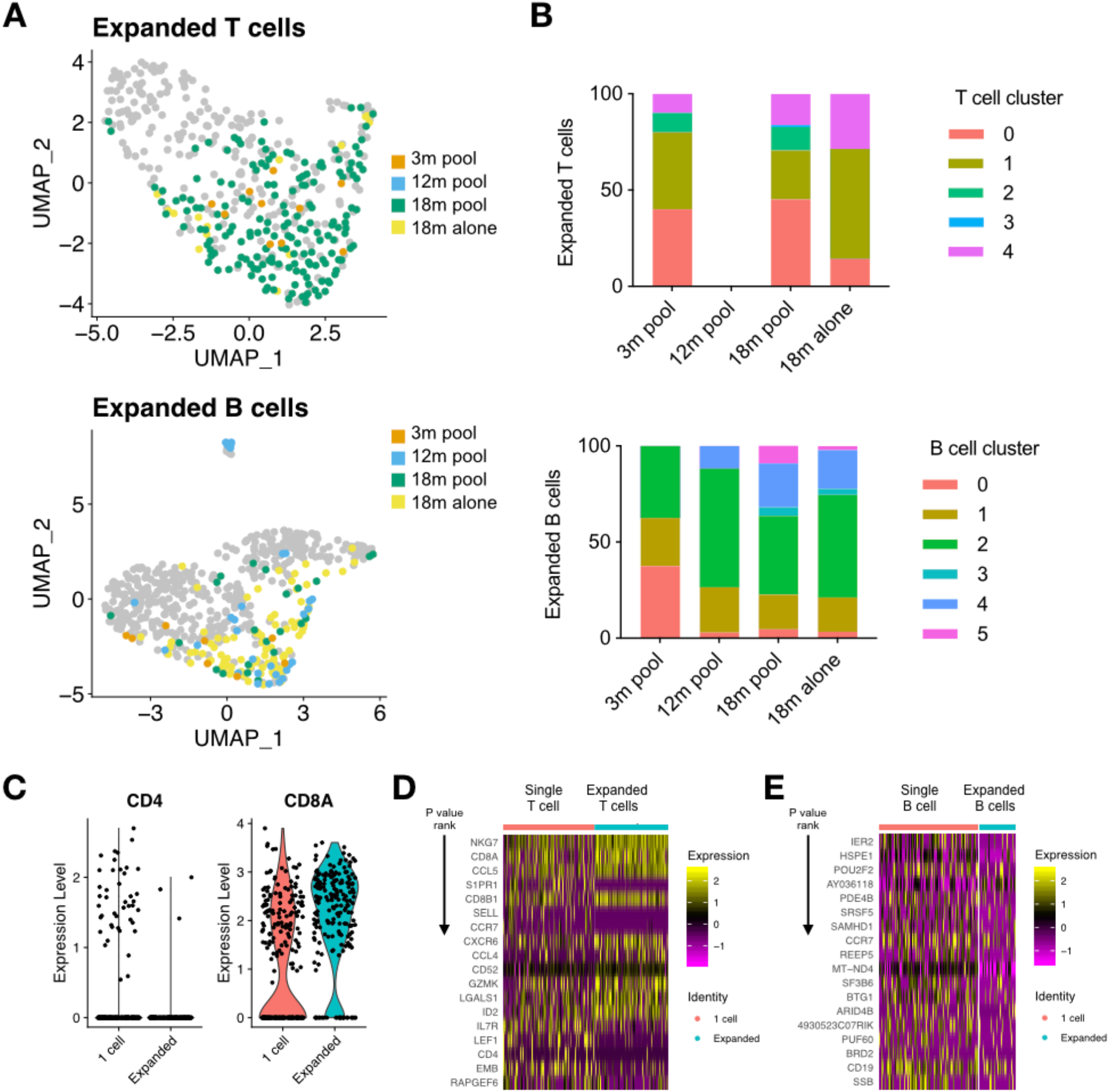
Clonally expanded lymphocytes have distinct transcriptional profiles. (A) Location and cluster membership of expanded B and T cells. Expanded B and T cells were defined by those clones containing at least 2 distinct cell barcodes. Clone is defined as identical, paired amino acid CDR3 sequences (CDRβ3+CDRα3 for T cells, CDRH3+CDRL3 for B cells) with exactly one heavy chain and one light chain. (B) Cluster membership of clonally expanded T and B cells in V(D)J libraries. Clonally expanded T and B cells correspond to those clones (identical, paired CDR3 amino acid sequence) containing two or more distinct cell barcodes. (C) CD4 and CD8a expression for those clones supported by a single cell or those expanded clones (more than 1 cell). (D-E) Heatmap of top differentially expressed genes between T cell (D) and B cell (E) clones with either one cell or clonally expanded. Gene order corresponds to adjusted p value. All genes displayed correspond to multiple hypothesis corrected p values less than 0.01. Heatmap intensity depicts normalized gene expression.

## Discussion

Here we present a comprehensive single-cell analysis of immune repertoires and transcriptomes of lymphocytes in the context of aging and CNS. Our experimental and computational workflow revealed the presence of clonally expanded B and T cells in the brains of aged mice. While previous studies have demonstrated increased frequencies of lymphocytes in the diseased CNS ^2^, it could not be determined whether these cells were clonally related. Our single-cell sequencing approach revealed that lymphocyte CNS infiltration was driven by clonally expanded lineages. Additionally, we were able to profile differences in gene expression between expanded and unexpanded lymphocytes. Further experiments combining immune repertoire sequencing from blood and secondary lymphoid organs could answer whether these expanded clones are specific to the CNS environment or expanded across different physiological compartments. Investigating whether identical clones would be present in the repertoires across various peripheral organs within the same aged individual would highlight whether the expansion we observed is a CNS-specific phenomenon, or if there is a global accumulation of clonally related lymphocytes in organs typically unassociated with immunity. The discovery of clonally expanded lymphocytes in the aged CNS suggests that determining their cognate antigens would be important in order to elucidate which role, if any, they play under homeostatic and disease states. Relating antigen-specificity information to immune receptor sequence could further inform whether these expanded B and T cell clones originate from cross-reactivity, dysregulation of self-tolerance, or simply past environmental challenges. Along these lines, determining whether environmental exposure (e.g. diet and hygiene) influences peripheral immune expansion would be of interest for future work.

Recently clinical data has indicated that therapeutic antibodies have demonstrated clinical value in treating neurological disorders, such as multiple sclerosis ^18,44^. Investigating whether or not clonally expanded lymphocytes are impacted by such treatments could be crucial to understanding and developing disease modifying treatments for neurological disorders for Alzheimer’s disease, multiple sclerosis or brain cancer. Expanded lymphocytes could represent a population of clones that persistently interact with their cognate antigen during homeostatic ageing, and following a disease or infection trigger, such cells may be poised to exacerbate immunopathology. In this case, the high number of predominantly IgM and IgD B cell clones might arise in an antigen-activated manner independent of T cell help. Alternatively, previous reports have demonstrated that an increase in IgD expression coincides with a decrease in IgM levels in the context of autoimmunity and anergy ^45,46^, providing one possible explanation to the high number of IgD-exclusive B cell clones found in aged CNS. Another hypothesis regarding the role of clonally expanded CNS lymphocytes arises from a recent study that identified virus-reactive T cells in the brains of Alzheimer’s patients and even provided an example of a cognate antigen ^2^. Future experiments performing a similar validation of clonally expanded B and T cells against pathogen or autoimmune targets could shed light upon age-associated immune dysregulation. In conclusion, our findings demonstrated the presence of clonally expanded lymphocytes in the CNS and furthermore quantitatively describe the immune repertoire and transcriptome landscape accompanying homeostatic ageing.

## Methods

### Murine brain cell isolation

Mouse experiments were performed under the guidelines and protocols approved by the Basel-Stadt cantonal veterinary office (Basel-Stadt Kantonales Veterinäramt Tierversuchsbewilligung #2582). 3-month-old, 12-month-old, and 18-month-old male C57/Bl6j mice were purchased (Janvier Laboratories, France) and were housed under pathogen-free conditions and maintained on a standard chow diet. Mice were housed for two weeks before sacrifice and brain extraction. Mice were perfused with cold PBS for 3 minutes following sacrifice and entire brains were collected in 1mL of RPMI. Brains were subsequently transferred to a new tube containing 1mL RPMI with Collagenase I/DNAse I (Roche) and cut to smaller pieces. Following 30-minute incubation at 37°C, each brain was mashed through an individual 70-micron cell strainer into a 50mL falcon tube. For three of the conditions, single-cell suspensions were pooled following meshing. Following a 30% Percoll gradient, cells were incubated with fluorescently conjugated antibodies for 30 minutes at 4°C. Antibodies used in the experiment: CD8a-BV421 (Biolegend, clone 53-6.7), CD4-PE (Biolegend, clone GK1.5), B220-APC (Biolegend, clone RA3-6B2), CD19-PE-Cy7 (Biolegend, clone 6D5), CD3e-FITC (Biolegend, clone 145-2C11), NK1.1-APC-Cy7 (Biolegend, clone PX136), Ter119-APC-Cy7 (Biolegend, clone TER-119). Multi-parametric flow cytometric sorting was performed using a FACSAria with FACSDiva software. B cells (TER-119^−^, NK1.1^−^, CD19^+^, B220^+^), CD4 T cells (TER-119^−^, NK1.1^−^, CD3e^+^, CD4^+^) and CD8 T (TER-119^−^, NK1.1^−^,CD3e^+^, CD8^+^) cells were sorted into separate 1.5mL eppendort tubes and pooled for the four age cohorts before proceeeding with sequencing library preparation.

### Single-cell sequencing libraries

Single cell 10X libraries were constructed from the isolated single cells following the Chromium Single Cell V(D)J Reagent Kits User Guide (CG000086 Rev M). Briefly, single cells were co-encapsulated with gel beads (10X Genomics, 1000006) in droplets using 4 lanes of one Chromium Single Cell A Chip (10X Genomics, 1000009). V(D)J library construction was carried out using the Chromium Single Cell 5’ Library Kit (10X Genomics, 1000006) and the Chromium Single Cell V(D)J Enrichment Kit, Mouse B Cell (10X Genomics, 1000072) and Mouse T Cell (10X Genomics, 1000071). The reverse transcribed cDNA was split in three and GEX, B and T cell V(D)J libraries were constructed following the instructions from the manufacturer. Final V(D)J libraries were pooled and sequenced on the Illumina NovaSeq platform (300 cycles, paired-end reads). Pooled 5’ gene expression libraries were and sequenced on the Illumina NextSeq 500 (26/91 cycles, paired-end) with a concentration of 1.6 pM with 1% PhiX.

### Repertoire analysis

Raw sequencing files arising from multiple Illumina sequencing lanes were merged and supplied as input to the command line program cellranger (v3.1.0) on a high-performance cluster. Raw reads were aligned to the germline segments from the GRCm38 reference and subsequently assigned into clonal families based on identical combinations of CDR3 (CDRH3+CDRL3 for B cells, CDRβ3+CDRα3 for T cells) amino acid sequences. Only those clonal families containing exactly one heavy chain and one light chain sequence were maintained in the analysis. Isotype was determined based on the constant region alignment for the majority of cells within each clonal family. In the case that the variable region alignment was provided but isotype was not recovered the isotype was labeled as “Undetermined”. Germline gene usage was determined by cellranger’s vdj alignment to the murine reference. Information involving UMI count, full-length sequences, germline gene annotations was extracted from the contig files produced by cellranger’s vdj pipeline. Network analysis was performed using the R package igraph ^47^. Edit distances were calculated using the stringdist package on the pasted CDRH3+CDRL3 sequences (or CDRβ+CDRα sequences for T cells) ^48^. Somatic variations were calculated by first extracting amino acid and nucleotide sequences of the entire VDJRegion (defined by the region spanning from framework region 1 on the V segment to framework region 4 on the J segment) based on initial alignment to the built in murine reference alleles and subsequent exportAlignments function of MiXCR v3.0.1 ^49^. The full-length, heavy and light VDJRegion sequences containing identical cellular barcodes were appended and then the number of clonal variants was determined by calculating the number of unique amino acid sequences. Somatic hypermutation was quantified for each heavy and light chain based on the number of nucleotide substitutions in the V and J regions compared to those germline genes with the highest alignment scores determined by MiXCR.

### RNA-Seq analysis

The output matrices, barcodes, and features files from cellranger (10X Genomics) for each sample were supplied as input to Seurat using the Read10X function and subsequently converted into a Seurat object using the function CreateSeuratObject. All BCR and TCR related genes (Ig V(D)J genes, isotype constant regions, J-chain, TRB V(D)J) were filtered out prior to further analyses. Data was normalized using a scaling factor of 10000 and variable features were found using 2000 genes. Cluster resolution was set to 0.5 and the first ten PCR dimensions were used for neighborhood detection. UMAP was performed again using the first ten PCA dimensions. All repertoires were proceeded and filtered together. Genes defining clusters were determined setting the min.pct argument equal to 0.25 using Seurat’s FindMarkers function. For analyses involving T and B cells found in repertoire sequencing data, only those cells containing identical nucleotide barcodes sequences were extracted and reanalyzed in the Seurat object. Heatmaps of cluster defining genes were selected for the top genes ranked by p value after employing the Bonferroni correction for multiple hypothesis testing.

## Acknowledgements

We acknowledge and thank Dr. Christian Beisel, Elodie Burcklen, Ina Nissan, and Tobias Schär at the ETH Zurich D-BSSE Genomics Facility Basel for excellent support and assistance. We also thank Mariangela Di Tacchio and Marie-Didiée Hussherr for excellent experimental support.

## Author Contributions

AY and DN performed experiments and analyses. All authors contributed to the study and manuscript design.

## Data and Materials Availability

The raw FASTQ files from deep sequencing that support the findings of this study will be deposited (following peer-review and publication) in the European Bioinformatics Institute. Additional data that support the findings of this study are available from the corresponding author upon reasonable request.

---

**Figure S1.**
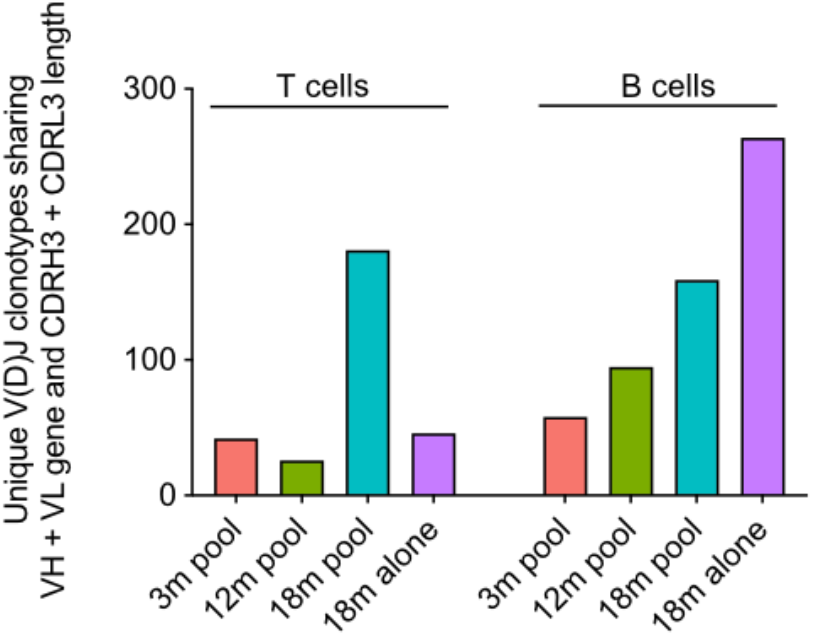
Relaxing clonotype definition minorly impacts clonal diversity. The number of clones for B and T cells are depicted after clustering those clones utilizing similar variable germline genes and containing identical CDR3 lengths (for both chains).

**Figure S2.**
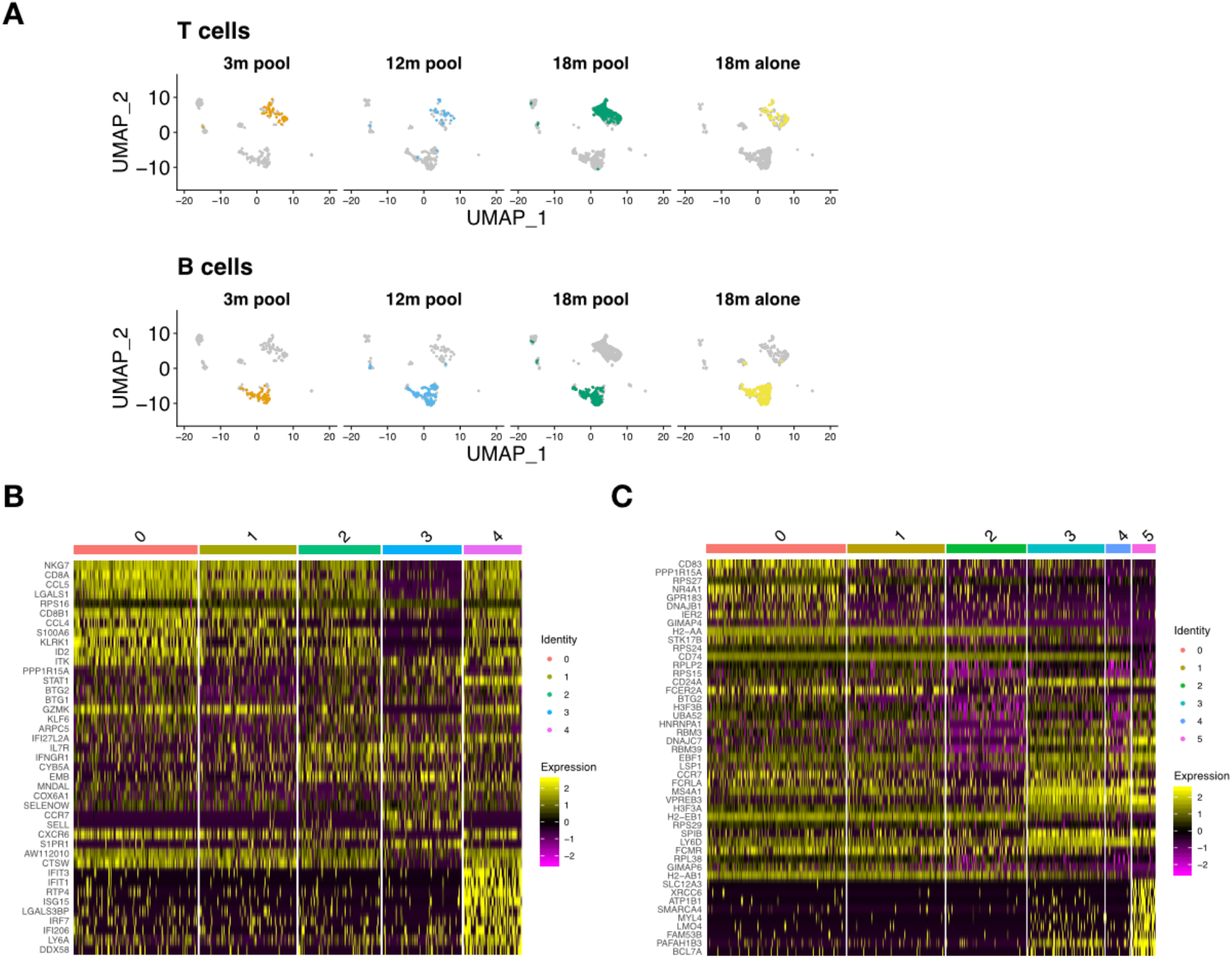
Transcriptional profiles of lymphocytes detected in V(D)J sequencing libraries. (A) Cells found in both gene expression and V(D)J sequencing libraries were highlighted on the uniform manifold approximation projection (UMAP) previously calculated using all recovered cells from gene expression libraries. Only those barcodes containing paired, full-length immune receptor information were highlighted. (B-C) Heatmap of the top 10 variable genes in each cluster for T and B cells found in V(D)J libraries, respectively. Intensity corresponds to normalized gene expression and row color indicates cluster membership.

**Figure S3.**
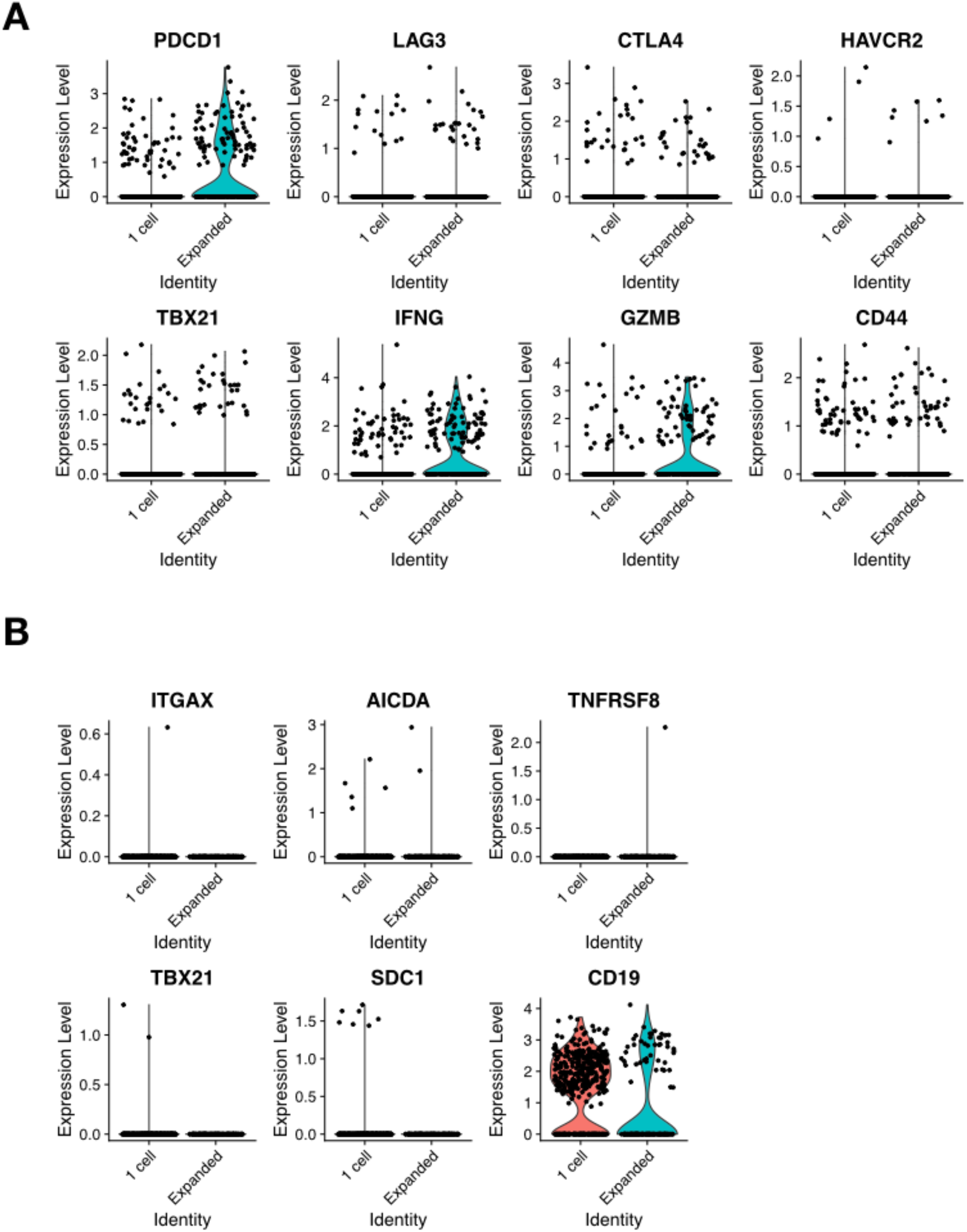
Expression levels of genes relevant to T and B cell function. (A-B) Normalized expression levels of select genes for T (A) and B (B) cells found in V(D)J libraries.

**Figure S4.**
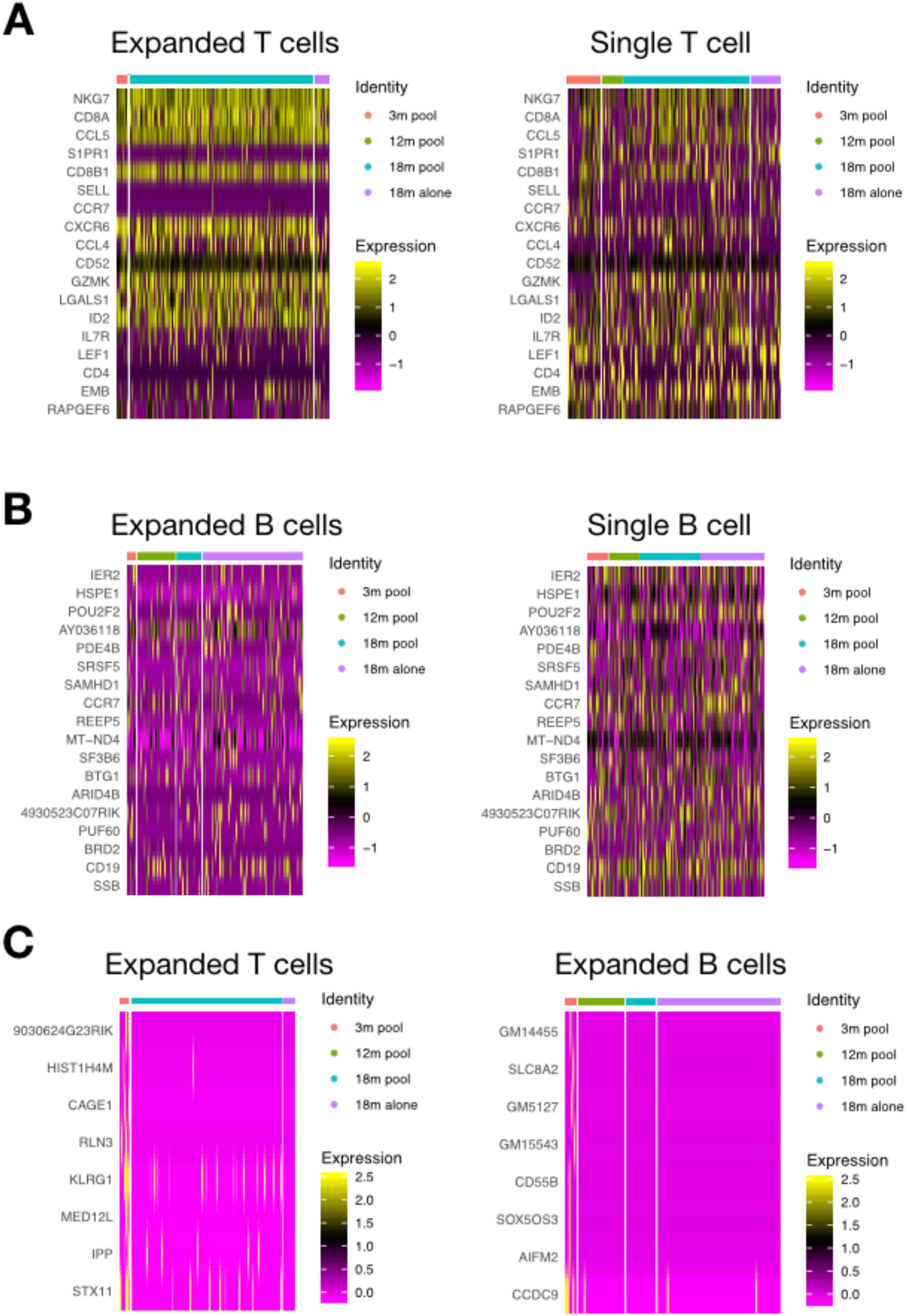
Minor transcriptional differences between CNS clonally expanded lymphocytes in young and old mice. (A-B) Heatmap of the top differentially expressed genes between expanded and unexpanded lymphocytes are ranked by adjusted p value. Intensity depicts normalized gene expression. (C) Heatmap of the top differentially expressed genes after comparing expanded T (left) and B (right) cells in 3-month-old cohort versus the three elderly cohorts. Intensity depicts normalized gene expression.

